# Sarcolemmal and mitochondrial membrane potentials measured *ex vivo* and *in vivo* in the heart by pharmacokinetic modelling of [^99m^Tc]sestamibi

**DOI:** 10.1101/2024.09.05.611418

**Authors:** Edward CT. Waters, Friedrich Baark, Matthew R. Orton, Michael J. Shattock, Richard Southworth, Thomas R. Eykyn

**Affiliations:** School of Biomedical Engineering and Imaging Sciences, King’s College London, The Rayne Institute, St. Thomas’ Hospital, London, SE1 7EH, UK; Department of Radiology, MRI Unit, The Royal Marsden NHS Foundation Trust, London, UK; Division of Radiotherapy and Imaging, The Institute of Cancer Research, London, UK; School of Cardiovascular and Metabolic Medicine and Sciences, King’s College London, United Kingdom

**Keywords:** [^99m^Tc]sestamibi pharmacokinetics, membrane potential, mitochondrial dysfunction, lipophilic cations, cardiac metabolism

## Abstract

We present a compartmental modelling approach to analyse radioactive time activity curves for first pass kinetics of [^99m^Tc]sestamibi in the heart. Reparametrizing the kinetic equations using the Nernst membrane-potential equation provides a novel means of non-invasively estimating the sarcolemmal (*E*_*m*_) and mitochondrial (ΔΨ_*m*_) membrane potentials in the heart. A Markov Chain Monte Carlo (MCMC) fitting approach was applied to data derived from established interventions in Langendorff perfused rat hearts where the sarcolemmal membrane was depolarised using hyperkalaemic Krebs Henseleit buffers; the mitochondrial membrane was depolarised using carbonylcyanide-3-chlorophenylhydrazone (CCCP); or both membranes were depolarised using their combination. Translating this approach to single photon emission planar scintigraphy kinetics from healthy rats allowed an estimate of these membrane potentials (voltages) *in vivo* for the first time; the values were *E*_*m*_ = –62 ± 5 mV and ΔΨ_*m*_ = –151 ± 5 mV (n = 4, mean ± SD).

## Introduction

Originally investigated by Hodgkin, Katz and Huxley^1,2^ and Mitchell,^3^ respectively, the transmembrane electrical potentials across the plasma and inner mitochondrial membranes are fundamental attributes of almost all mammalian cells. Because of their importance in normal cellular function, estimating membrane potentials *in vivo* is needed to understand cellular physiology and pathophysiology; yet it continues to be a major technical challenge. Mitochondrial dysfunction is an integral part of metabolic dysregulation, and it underlies many diseases including (cardio)myopathy, diabetes, chronic inflammation, degenerative diseases, and cancer.^4^

In excitable cells, the resting membrane potential, *E*_*m*_ is typically maintained at –50 to –80 mV. It is a fluctuating quantity that is modulated by the plasma membrane’s ‘action potential’ in smooth and skeletal muscle fibres, the heart, and neurons. In many excitable cells including cardiac myocytes, the action potential is initiated by the opening of voltage-gated Na^+^ channels, permeability to Na^+^ increases, leading to electrical depolarization to about +40 mV. The internal membrane potential can be measured in isolated cells by voltage clamping the plasma membrane potential. Sarcolemmal membrane potential can also be measured in the arrested perfused hearts using KCl containing micro electrodes.^5^ *E*_*m*_ is given by the ratio of the equilibrium concentrations of the major permeant monovalent ions K^+^, Na^+^, and Cl^-^, inside and outside the cell, and their relative permeabilities. These are included in the Goldman-Hodgkin-Katz (GHK) equation that is used to calculate the membrane potential. The membrane potential is reported to be depolarized in proliferating cells and some cancer cells,^6^ and it plays a major role in systolic and diastolic dysfunction in cardiovascular pathologies such as ischaemia, heart failure and arrhythmias.

In contrast, the resting mitochondrial membrane potential ΔΨ_*m*_ is tightly controlled between –135 to –165 mV,^7^ but may become depolarized or hyperpolarized under stress or pathophysiological conditions. The transmembrane voltage difference achieved by the transport of electrons along the electron transport chain, and the pumping of H^+^ from the mitochondrial matrix into the inter membrane space, creates an electrochemical gradient across the inner mitochondrial membrane with a pH gradient (Δ*pH* typically about 1 pH unit higher in the mitochondrial matrix) that collectively contributes to the proton motive force (Δp) that drives ATP synthesis via the F_0_F_1_-ATPase.^3^ Methods for measuring mitochondrial membrane potential typically use lipophilic cationic molecules^8,9^ that passively diffuse across the membrane; *e*.*g*., tetramethylrhodamine ethyl ester (TMRE), ethidium, tetraphenylphosphonium (TPP), triphenylmethylphosphonium (TPMP), and tetraphenylarsonium (TPA).^7^ The subcellular distribution of such tracers at equilibrium can be used to estimate the trans-membrane potential (voltage) using the Nernst equation; this analysis assumes that probe molecules do not significantly perturb the membrane potential, which they are designed to measure. ΔΨ_*m*_ has been estimated in perfused rat hearts with [^3^H]triphenylmethylphosphonium to be –125 ± 7 mV at the normal physiological heart rate of 300 beats per minute (bpm), becoming hyperpolarized under low energy demand^10^ or depolarized under high energy demand.^11^ Such experimental approaches require sub-cellular fractionation to measure concentrations of the probe molecules in the different cellular compartments; thus they are intrinsically destructive of the tissue, *viz*., invasive.

Non-invasive measurement of membrane potentials *in vivo* using radiolabeled lipophilic cations with positron emission tomography (PET) have been reported.^12-16^ Total membrane potential *E*_*t*_ = *E*_*m*_ + ΔΨ_*m*_ (the sum of both sarcolemmal and mitochondrial membrane potentials) can be measured from the steady state distribution of [^11^C]TPMP,^12^ [^18^F]TPP,^13^ or [4-^18^F]fluorobenzyl-triphenylphosphonium (^18^F-BnTP),^14^ while contrast-enhanced MRI using gadolinium can be used to measure extracellular volume fractions.^15,16^ [^99m^Tc]sestamibi, and related tracers, such as [^99m^Tc]tetrofosmin are single photon emission computed tomography (SPECT) agents that are lipophilic cations, and therefore their cellular distribution and tissue retention is governed by the negative potential across both sets of membranes.^17^ Sestamibi uptake is sensitive to uncoupling oxidative phosphorylation in mitochondria with m-chlorophenylhydrazone (CCCP), with a sensitivity that is comparable to that of TMRE;^17^ and to doxorubicin-induced cardiotoxicity, which is independent of cardiac perfusion.^18^

Thus, in view of the shortcomings with previous dye-distribution methodologies for in vivo imaging we aimed to develop a new approach that uses the Nernst equation in a kinetic (rate equation) model to fit numerically to experimental time activity data of [^99m^Tc]sestamibi. The model equations describe the rates of [^99m^Tc]sestamibi uptake in isolated Langendorff-perfused hearts *ex vivo* under control conditions, and when either (or both) membranes are depolarized pharmacologically. The method was evaluated by using a least squares fitting method combined with a Markov Chain Monte Carlo (MCMC) method to estimate parameters by systematic random sampling of high-dimensional probability distributions.^19^ We applied our approach to high temporal resolution single photon emission planar scintigraphy imaging data yielding, for the first time, non-destructive estimates of the two fundamental electrical-potential differences in the heart *in vivo*.

## Theory

### Goldman-Hodgkin-Katz (GHK) and Nernst equations

The sarcolemmal membrane potential (*E*_*m*_ in Volts) is given by the ratio of the concentrations at equilibrium of the major permeant ions K^+^, Na^+^, and Cl^-^, inside and outside the cell, given by the Goldman-Hodgkin-Katz (GHK) equation,

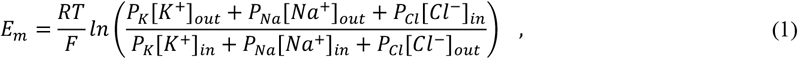

Simulations of the sarcolemmal membrane potential were performed by calculating *E*_*m*_ over the range of [*K*^+^]_*out*_ values for the different experimental KHB buffers and fixing the other parameter values. Typical concentrations in cardiac tissue are (mM) [*K*^+^]_*in*_ = 113, [*K*^+^]_*out*_ = 5.9, [*Na*^+^]_*out*_ = 140, [*Na*^+^]_*in*_ = 10, [*Cl*^−^]_*out*_ = 140 and [*Cl*^−^]_*in*_ = 10, with *relative* permeabilities of *P*_*K*_ = 1, *P*_*Cl*_ = 0.5 and *P*_*Na*_ = 0.02.^20^ The Nernst equation for the [*K*^+^] equilibrium (*E*_*K*_) can be simulated by setting *P*_*Na*_ = *P*_*Cl*_ = 0 in Eq. 1.

### Three-compartment model describing ^99m^Tc sestamibi pharmacokinetics

The compartmental model shown in Fig. 1A describes the delivery of an arterial bolus input *U*(*t*) to give a plasma compartment *c*_*p*_(*t*) which can leave the tissue via the venous outflow or diffuse across the sarcolemmal membrane at a rate characterized by the value of the rate constant *k*_1_. Molecules in the cytosolic compartment *c*_*t*_(*t*) can diffuse back out of the cell at a rate characterized by the value of the rate constants constant *k*_–1_, or they can diffuse into or out of the mitochondria at rates characterized by the value of the rate constants *k*_2_ and *k*_–2_, respectively, to give a mitochondrial compartment *c*_*m*_(*t*).

**Figure 1.**
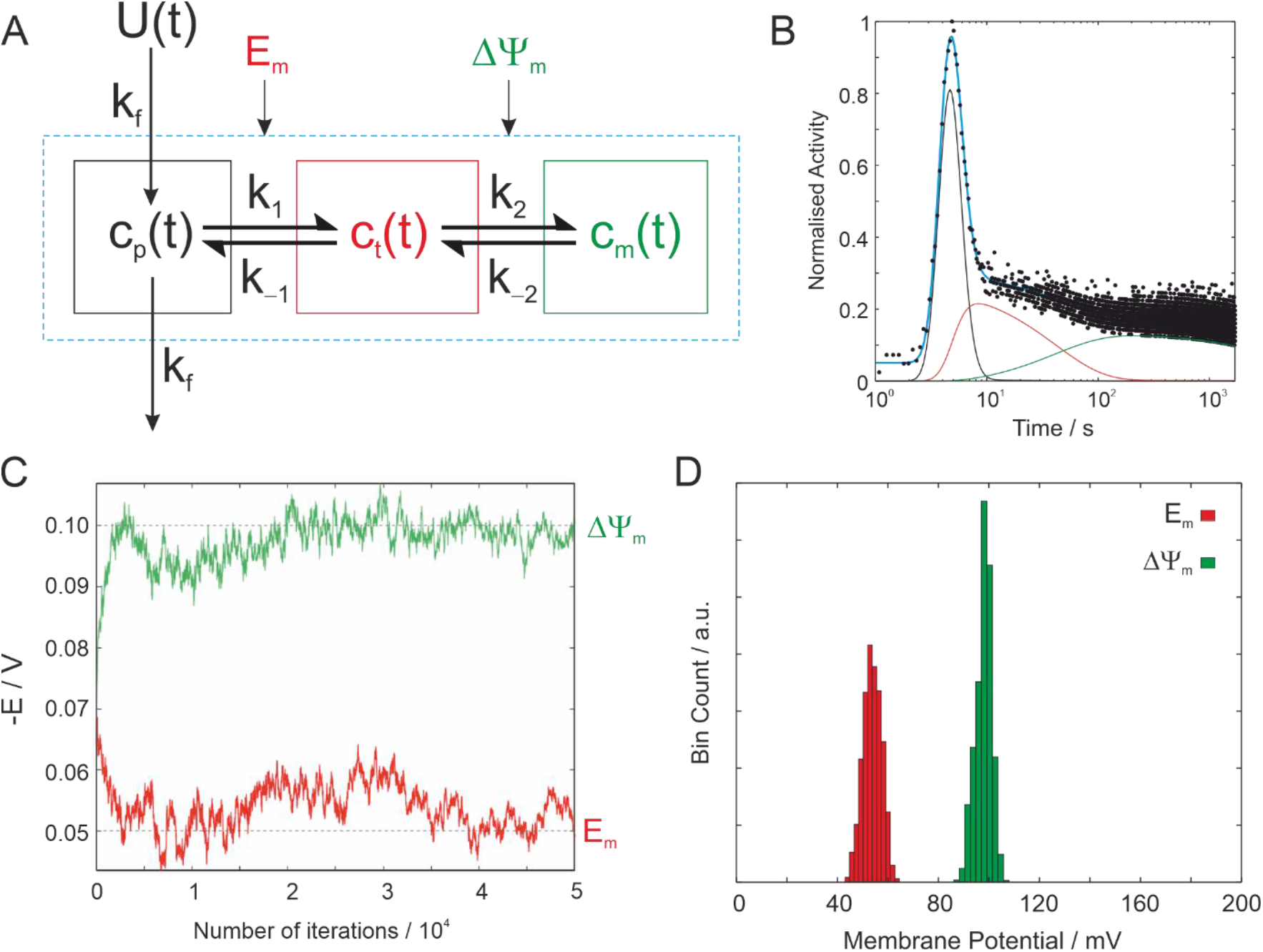
**A** Three compartment model describing the passage of a radiotracer through the heart where *c*_*p*_(*t*) (black), *c*_*t*_(*t*) (red) and *c*_*m*_(*t*) (green) are the plasma, cytosolic and mitochondrial concentrations, respectively. The bolus input function is given by U(t), which is delivered into or out of the plasma compartment by flow described by the rate constant k_f_. The rate constants across the sarcolemmal membrane are k_1_ and k_-1_. The rate constants across the mitochondrial membrane are k_2_ and k_-2_. The sarcolemmal and mitochondrial membrane potentials are given by the voltages E_m_ and ΔΨ_*m*_, respectively. **B** Simulated data with known (true) parameters of *E*_*m*_ = –50 mV and ΔΨ_*m*_ = –100 mV showing the separate compartments with black, red and green solid lines. **C** Output of the Markov Chain Monte Carlo random walk for the simulated data in **B**. The true parameters are denoted by the dashed lines. For illustrative purposes the initial input parameters in the MCMC were set to *E*_*m*_ = –65 mV and ΔΨ_*m*_ = –70 mV. An initial burn in period can be seen as the algorithm establishes a stable equilibrium close to the true values of the simulated parameters. The solid cyan line in **B** is the result of the MCMC fit. **D** Histogram showing the distribution of membrane potentials from the MCMC fit from which means and standard deviations can be calculated.

Assuming ionic solutes are present at tracer concentration, and the membrane potential is not perturbed by their redistribution across the membranes, then the ratio of the concentrations across the membrane at steady state depends on the membrane potential described be the Nernst equation. From standard chemical kinetics, the equilibrium constant is given by the ratio of the rate constants *k*_1_/*k*_–1_ and *k*_2_/*k*_–2_ even for a system not at a steady state. The reverse rate constants *k*_–1_ and *k*_–2_ can therefore be expressed as a function of the forwards rate constants and the corresponding membrane potentials. The system of differential equations describing the model in Fig 1 is expressed as a function of concentrations and membrane potentials as:

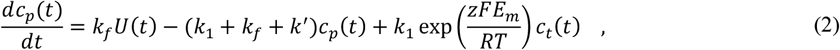

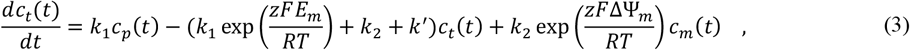

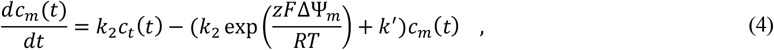

Where the universal gas constant is R = 8.3144621 J mol^-1^ K^-1^, the Faraday constant F = 96485 C mol^-1^ (kJ V^-1^ mol^-1^), the temperature T = 310 K, and *z* is the ion’s charge; *E*_*m*_ and ΔΨ_*m*_ are the sarcolemmal and mitochondrial membrane potentials, respectively, which are defined as negative voltages in the above equations. The first order rate constant *k*^′^ = 3.2059 × 10^−5^ s^−1^ describes the radioactive decay rate constant of ^99m^Tc (*k*^′^ = ln 2 /*λ* where *λ* = 6.006 h is the half-life of ^99m^Tc). Radioactivity is measured in counts per second (cps) and is not a direct measure of concentration *per se*. However, activity per unit compartment volume is proportional to concentration (mol per unit volume). Providing the kinetic model is first order, as is the case here, then the rate constants *k* (units s^-1^) are independent of concentrations.

The bolus input function is represented by a modified gamma-variate function given by:

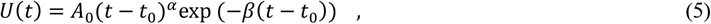

where *t*_0_ is the initial arrival time of the bolus. *A*_0_ is a normalization factor, and *α* and *β* characterise the rise and fall of the concentration of the input bolus, respectively, *i*.*e*., the width or dispersion of the bolus. The measured signal (radioactivity) is the volume weighted average of the three compartments, given by:

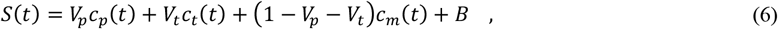

where *B* is a fitting parameter that allows for non-zero baseline offset due to Poisson noise that arises from detecting radioactive counts.

For first-pass flux of a solute into the tissue the rate constant *k*_1_ depends on blood flow and extraction fraction, with expressions derived by Renkin and Cone.^21,22^ If the membrane permeability is low and/or the surface area of the capillary membrane is small compared to flow, then the rate constant is dependent on membrane transport and independent of flow. Membrane permeability of [^99m^Tc]sestamibi is lower than flow tracers and is independent of flow at high perfusion rates, becoming flow dependent when perfusion decreases, for example during ischaemia.^23^ Normal coronary arterial perfusion in the human heart is 2.4 ml g^-1^ min^-1^,^24^ with slightly higher values in the rat heart *in vivo*.^25^ The rate of arterial perfusion in the Langendorff perfused rat heart is much higher at 14 ml g^-1^ min^-1^. When perfusion is not rate limiting, then the rate constant *k*_1_ is dependent on diffusion mediated transport, which depends on the membrane potential (voltage).

## Results

A Markov Chain Monte Carlo (MCMC) method was employed to estimate the model parameters by systematic random sampling of high-dimensional probability distributions. Time activity curves were simulated using Eqs. (2-6) with the same temporal resolution as the experimental data and known (true) parameters to explore the robustness of the fitting and associated errors. The input function was modelled as a gamma variate function with *A*_0_ = 1, *α* = 12, *β* = 4 assuming zero signal at *t* < *t*_0_. Rate constants were set to *k*_*f*_ = 0.5, *k*_1_ = 0.1 and *k*_2_ = 0.01 and a range of membrane potentials. Fig. 1B shows a simulated dataset where *E*_*m*_ = –50 mV and ΔΨ_*m*_ = –100 mV. Randomly distributed Poisson noise was added and the fitting parameter *B* was set to zero for simulated curves. Fig. 1C shows a typical MCMC random walk for the simulated data. There is an initial burn in period which allows the algorithm to achieve a stable equilibrium after which it converges on the known true parameter values shown by dashed lines. The resulting fit from the MCMC procedure is shown as a solid cyan line in Fig 1B and a histogram of fit parameters in Fig 1D from which the mean and standard deviation were calculated.

The method was applied to fitting experimental time activity curves measured following bolus injection of [^99m^Tc]sestamibi in the perfused rat heart with perfusion maintained at 14 ml min^-1^ in all hearts for the duration of each experiment. Experiments and MCMC fitting were performed with a range of hyperkalaemic buffers with [K^+^] = 4.9, 10, 15, 20 and 25 mM. Fig. 2A shows a representative fit for a heart perfused with [K^+^] = 4.9 mM (i.e. control group) with the MCMC fit shown with a cyan line. The separate compartments in the fit are denoted by the solid black, red and green lines. Fig. 2B shows a histogram of the rates derived from 10,000 iterations of the MCMC algorithm. Sarcolemmal and mitochondrial membrane potentials in control hearts were estimated to be *E*_*m*_ = –64 ± 1 mV and ΔΨ_*m*_ = –101 ± 5 mV (n = 4, mean ± SD), respectively. Fig. 2C shows a representative fit for a heart perfused with high [K^+^] = 25 mM with the resulting MCMC histogram in Fig. 2D. The sarcolemmal membrane potential is shifted to a lower voltage in Fig. 2D compared to Fig. 2B. Cardiac contraction ceased when [K^+^] > 10 mM (Fig. 2E). Fig. 2F shows the mean values of *E*_*m*_ and ΔΨ_*m*_ for hearts perfused over the range of hyperkalaemic buffers. The sarcolemmal membrane was depolarized in a dose dependent manner with [K^+^] while the mitochondrial membrane potential was maintained over the 30 mins of perfusion. Fig. 2G shows *E*_*m*_ plotted as a function of [K^+^] showing a good agreement with the GHK equation (solid black line) compared to the simple Nernst equation (dashed black line).

**Figure 2.**
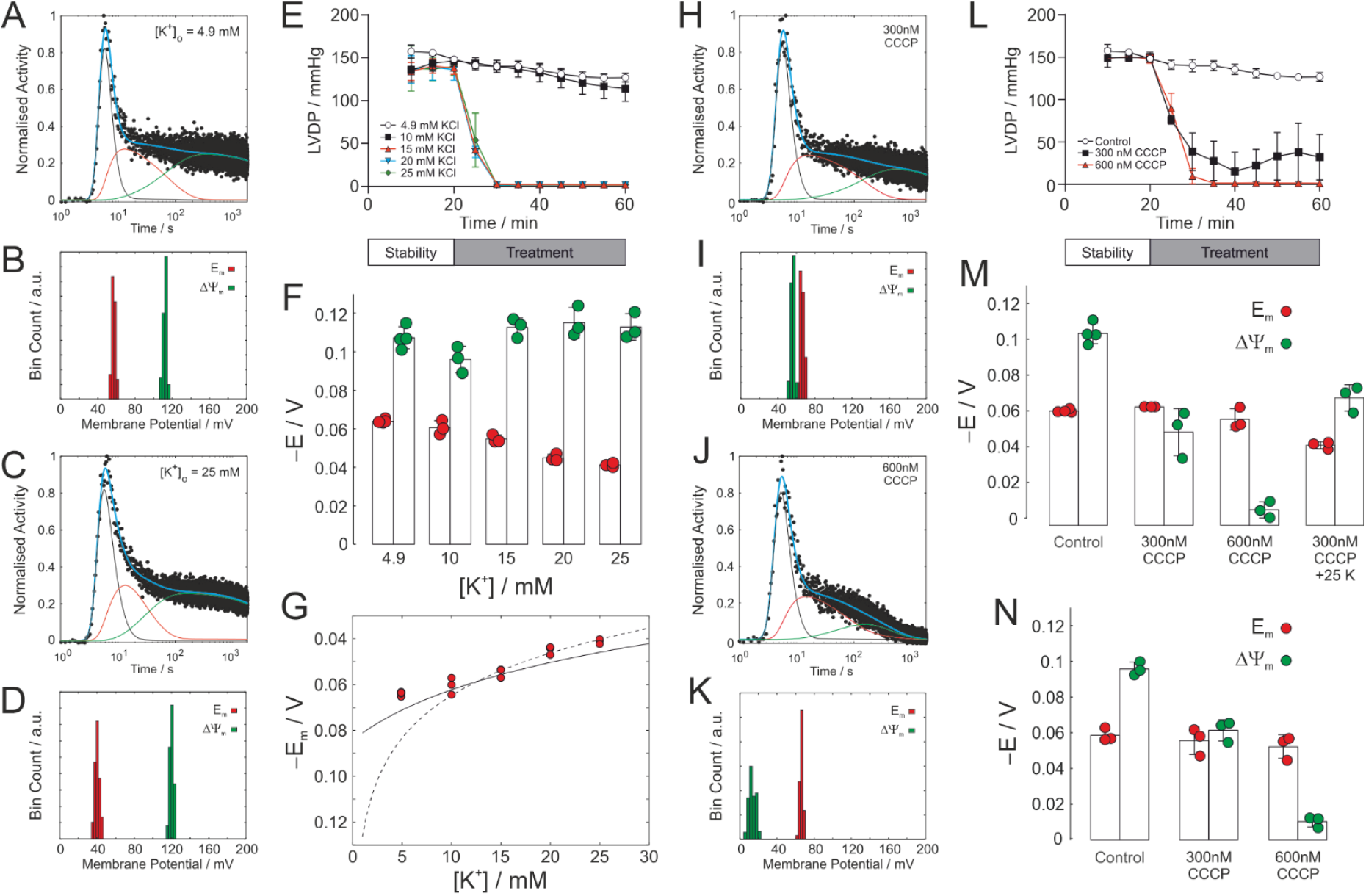
**A** Representative time activity curve for bolus injection of [^99m^Tc]sestamibi into a rat heart perfused with KHB containing 4.9 mM K^+^ and **B** a histogram of the derived rates resulting from 10,000 iterations of the MCMC algorithm. The cyan line shows the result of the best fit to the model. The three compartments *c*_*p*_(*t*), *c*_*t*_(*t*) and *c*_*m*_(*t*) are also shown in solid black, red and green lines, respectively. **C** Representative time activity curve for bolus injection of [^99m^Tc]sestamibi into a rat heart perfused with KHB containing 25 mM K^+^ and **D** a histogram of the derived rates resulting from MCMC analysis. **E** Left ventricular developed pressure (LVDP) measured for hearts perfused with control KHB and a range of hyperkalaemic buffers. **F** Plot of the mean values of *E*_*m*_ and ΔΨ_*m*_ derived from MCMC fitting of control KHB (n = 4) and the various hyperkalaemic KHB buffers (n = 3). **G** Values of *E*_*m*_ in panel **F** replotted with theoretical curves corresponding to the GHK equation (solid black line) and the Nernst equation (dashed line). **H** Representative time activity curve for bolus injection of [^99m^Tc]sestamibi into a rat heart perfused with KHB containing 300 nM CCCP and **I** a histogram of the derived rates resulting from MCMC analysis. **J** Representative time activity curve for bolus injection of [^99m^Tc]sestamibi into a rat heart perfused with KHB containing 600 nM CCCP and **K** a histogram of the derived rates resulting from MCMC analysis. **L** LVDP measured for hearts perfused with control KHB and 300 nM and 600 nM CCCP. **M** Plot of the mean values of *E*_*m*_ and ΔΨ_*m*_ derived from MCMC fitting of [^99m^Tc]sestamibi data for control KHB (n = 4), 300nM CCCP (n = 3), 600 nM CCCP (n = 3) and 300 nM CCCP + 25 mM K^+^ (n = 3). **N** Plot of the mean values of *E*_*m*_ and ΔΨ_*m*_ derived from MCMC fitting of [^99m^Tc]tetrofosmin data for control KHB (n = 3), 300nM CCCP (n = 3) and 600 nM CCCP (n = 3). All data are reported as mean ± SD.

Fig. 2H shows a representative fit for a heart perfused with the ionophore 300 nM carbonylcyanide-3-chlorophenylhydrazone (CCCP) alongside the resulting MCMC histogram (Fig. 2I) of the derived rates. Figs. 2J-K show the same for treatment with 600 nM CCCP. The measured mitochondrial membrane potential is shifted to a lower voltage in Fig. 2I compared to control hearts in Fig. 2B, and further still in Fig. 2K. Cardiac contraction, Fig. 2L, was significantly attenuated with 300 nM CCCP and arrested with 600 nM CCCP while perfusion was maintained at 14 ml min^-1^ for all hearts. Fig. 2M shows the mean values of *E*_*m*_ and ΔΨ_*m*_ for hearts perfused with 300 and 600 nM CCCP, which led to a depolarization of the mitochondrial membrane to ΔΨ_*m*_ = –52 ± 11 mV (n = 3, mean ± SD) and –9 ± 4 mV (n = 3, mean ± SD), respectively, but not the sarcolemmal membrane potential which was maintained over the 30 mins of perfusion. Both membrane potentials were depolarized, Fig. 2M, when treated with 300 nM CCCP + 25 mM [K^+^]. Time activity curves measured with [^99m^Tc]tetrofosmin were largely identical to those measured with sestamibi (data not shown). Fig. 2N shows the mean values of *E*_*m*_ and ΔΨ_*m*_ measured with tetrofosmin in control, 300 nM and 600 nM CCCP treated hearts, which were effectively identical to those measured with sestamibi.

Fig. 3A shows a time series of dynamic planar scintigraphy data of an anaesthetised rat and a region of interest positioned over the heart with a static planar image acquired 2 h post injection shown in Fig. 3B. The resultant time activity curve (Fig. 3C) differs from those measured in the perfused heart; the plasma compartment (which includes the ventricles in vivo) is narrower due to slower flow rates in vivo. Fig. 3D shows a histogram of the MCMC result. Fig. 3E shows the mean values of the sarcolemmal and mitochondrial membrane potentials measured in vivo in the rat heart which were *E*_*m*_ = –62 ± 5 *mV* and ΔΨ_*m*_ = –151 ± 5 *mV* (n = 4, mean ± SD), respectively.

**Figure 3.**
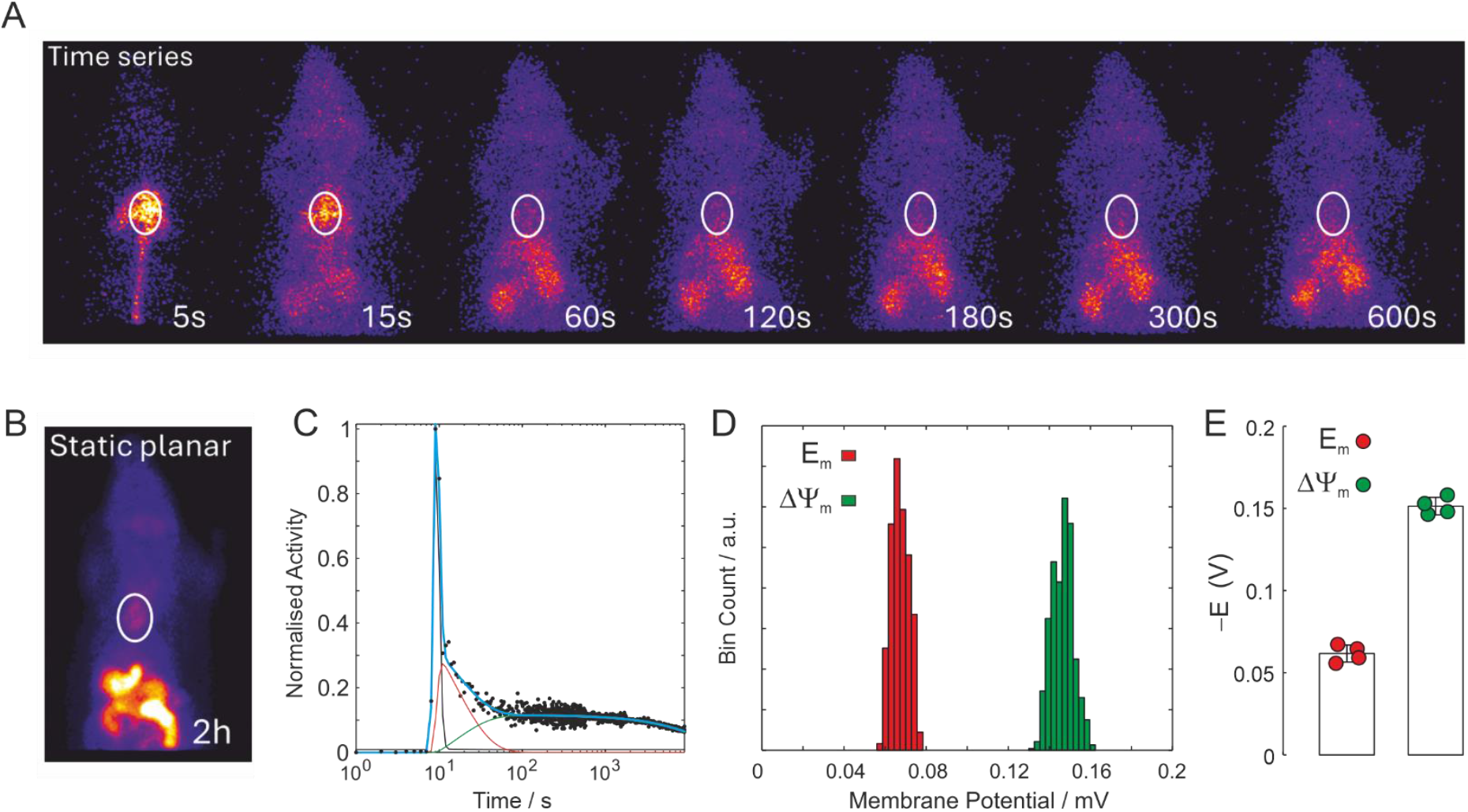
**A** Dynamic time series of SPECT planar scintigraphy images following bolus injection of [^99m^Tc]sestamibi in a healthy anaesthetised rat, and a region of interest positioned over the heart. **B** SPECT static planar scintigraphy image acquired 2 h post injection. **C** Representative time activity curve measured for the heart region of interest. **D** Histogram of the derived rates resulting from 10,000 iterations of the MCMC algorithm. **E** Sarcolemmal and mitochondrial membrane potentials measured *in vivo* in the rat heart were *E*_*m*_ = –62 ± 5 mV and ΔΨ_*m*_ = –151 ± 5 mV, respectively, (n = 4). Data are reported as mean ± SD.

## Discussion

Modelling the first pass kinetics of [^99m^Tc]sestamibi allowed an estimation of both sarcolemmal and mitochondrial membrane potentials in the heart ex vivo and notably in vivo. Titration with hyperkalaemic buffers led to a dose dependent decrease in the estimated *E*_*m*_ with increasing [K^+^]; this agreed well with the prediction made using the Goldman-Hodgkin-Katz equation. The sarcolemmal membrane potential *E*_*m*_ measured in vivo was the same as that measured in the perfused heart ex vivo. However, the magnitude was slightly lower than resting potentials previously reported.^5^ However, our measurements in control hearts ex vivo and in vivo reflect measurements that are time averaged over the entire cardiac cycle, with the diastolic membrane potential being more negative than our measurements and the systolic membrane potential being positive. It is of interest that the sarcolemmal membrane potential remained polarized during CCCP-mediated mitochondrial depolarization and cessation of cardiac contraction, reflecting the independence of sarcolemmal ion pumps on mitochondrial generation of ATP.^26-28^ Within the 30 minutes of these experiments, in the presence of maintained perfusion with a polarizing solution of glucose containing KHB, the sarcolemmal membrane does not depolarize. However, eventually, intracellular ATP will fall, K_ATP_ channels will activate and sarcolemmal ATPases would become inhibited. The mitochondrial membrane potential ΔΨ_*m*_ in the Langendorff perfused heart was slightly less negative than those reported in the literature^10^ and was found to be substantially more negative in vivo; this may reflect the suboptimal metabolism of a glucose-only perfused heart which lacks the full range of substrates and is likely less well oxygenated than when supplied with blood *in vivo*.

Previous studies have reported total myocardial membrane potential are *E*_*t*_ = –148.1 ± 6 mV in dogs, –146.7 ± 3.8 mV in rats and –139.3.1 ± 5.8 mV in mouse using [^11^C]TPMP^12^ while *E*_*t*_ = –160.7 ± 3.7 mV was reported in the human using [^18^F]TPP.^16^ These measurements were performed by continuous infusion of tracer which was allowed to reach a steady state, with the tracer distribution modelled as a two compartment system where the tissue compartment was subject to a single voltage *E*_*t*_ = *E*_*m*_ + ΔΨ_*m*_. The extracellular volume fraction was measured by contrast enhance MRI with blood sampling to measure plasma radioactivity. Our measurements gave *E*_*t*_ = –213 mV which does *not* agree with these previous measurements. This can be rationalized by noting that the previously reported values of *E*_*t*_ are in much better agreement with our values of ΔΨ_*m*_. Voltages of –62 mV and –151 mV gives equilibrium constants of *K*_1_ = 10 across the sarcolemmal membrane and *K*_2_ = 285 across the mitochondrial membrane. At these voltages the relative steady state concentrations in the plasma:cytosol:mitochondria are 1:10:2850, respectively; the plasma and cytosolic concentrations are negligible compared to mitochondria. Thus, it is possible that previously reported measurements of *E*_*t*_ were more representative of ΔΨ_*m*_.

While our new approach explicitly/directly yields estimates of *E*_*m*_ and ΔΨ_*m*_, possible (generic) limitations are associated with the complexity of the MCMC fitting method; and possible correlation of the fitted variables can arise. As is usual with this methodology, cross-correlation was assessed on simulated data (using the model of the system) by calculating a correlation matrix, coefficients of variance, and the fitting Jacobian. Fixing the values of some of the parameters, which might be estimated in separate experiments, reduces the variances of the fitted values. Another challenge arose in our analysis due to the large size of the data files with their very high temporal resolution slowing down the parameter fitting by the MCMC method. Such high temporal resolution was achieved by using gamma detectors and planar scintigraphy which is not possible with 3D SPECT imaging due to rotation of the camera.

The kinetic/compartment modelling approaches that we describe here, coupled with MCMC parameter estimation, can be readily applied to data acquired with SPECT imaging agents already in widespread clinical use. They could also potentially be translated to positron emitting lipophilic cations to harness the native capacity of PET for 3D dynamic scanning and deliver tomographic 3D images of mitochondrial and plasma membrane potentials. Recent advances in next-generation total body PET scanner technologies^29,30^ promise significant gains in sensitivity and temporal and spatial resolution, alongside the capacity for whole-body imaging, meaning that these approaches could be achieved with significantly lower radiation doses to patients. These approaches could provide a sensitive, calibratable and readily translatable new biomarker in patients, with many interesting new opportunities for the non-invasive study of whole-body physiology and disease in cardiovascular medicine and beyond.

## Materials and Methods

### Reagents and Gas Mixtures

All reagents were purchased from Sigma-Aldrich unless otherwise stated. All gas mixtures were purchased from BOC Industrial Gases.

### Animals

Male Wistar rats (280–320 g, Charles River, UK) were used for all experiments. All experimental procedures were approved by King’s College London’s local Animal Care and Ethics Committee and carried out in accordance with Home Office regulations as detailed in the Guidance on the Operation of Animals (Scientific Procedures) Act 1986.

### Heart Excision and Perfusion

Adult male Wistar rats (275–325 g) were used for all perfusion experiments. Rats were co-administered sodium pentobarbital (200 mg kg^−1^) and sodium heparin (200 IU kg^−1^) by intraperitoneal injection. Hearts were excised and immediately arrested in ice-cold Krebs–Henseleit buffer (KHB) consisting of (mmol L^-1^): NaCl, 118; KCl 5.9; MgSO_4_, 1.16; NaHCO_3_, 25; NaEDTA, 0.48; glucose, 11.1; and CaCl_2_, 2.2. The aorta was cannulated and secured using 3–0 suture (Ethicon), the pulmonary artery incised to drain coronary effluent, and perfused with KHB gassed with 95%O_2_/5%CO_2_ at 37°C, with perfusion maintained at a constant rate of 14 ml min^−1^. Contractile function was monitored with an intraventricular balloon set to an initial end-diastolic pressure of 4–10 mm Hg. Perfusion pressure and cardiac contractile function were recorded using two pressure transducers connected to a PowerLab data acquisition system (AD instruments Ltd). After a stabilisation period, perfusate supply was switched to either the vehicle control (0.02% v/v EtOH), 300-600 nM carbonylcyanide-3-chlorophenylhydrazone (CCCP) or 4.9-25 mM KCl in KHB. After 10 min of treatment a single bolus of either [^99m^Tc]sestamibi or [^99m^Tc]tetrofosmin (5 MBq, 50 μL), was introduced via the injection port.

### The Triple γ-Detector System

Radiotracer pharmacokinetics were monitored using a custom built triple γ-detector system described previously.^31,32^ This system consists of three orthogonally positioned lead-collimated Na/I γ-detectors arrayed around a Langendorff isolated heart perfusion apparatus. The detectors are sited: (i) 3 cm downstream of the injection port that was 15 cm upstream of the heart cannula on the arterial line; (ii) directly opposite the heart; and (iii) over the venous outflow tube. Each detector was connected to a modified GinaSTAR™ ITLC system running Gina™ software for real-time data collection (Raytest Ltd, UK). Cardiac accumulation of the tracer was measured with a temporal resolution of 200 ms and a total duration of 30 min.

### In vivo planar scintigraphy

Animals were anaesthetised in an induction chamber using isoflurane, 4% v/v at a flowrate of 0.5-1 L min^-1^. They were then maintained on a nose cone at 2% v/v isoflurane at a flowrate of 0.5-1 L min^-1^ and taped down in a supine position on a heating pad (MouseMonitor S, Indus) to maintain body temperature at 37°C, as measured by a rectal probe (MLT1403, AD Instruments). All nuclear imaging experiments were conducted using a Nanoscan SPECT/CT (Mediso, Hungary) scanner. Once animals had been transferred to the scanner, radiotracer was administered (1 ml of 100 MBq in sterile filtered saline) intravenously through the femoral vein. Scanning was initiated just prior to radiotracer injection to visualise the injection spike as the radiotracer enters the animal, to accurately sample the complete time course. Planar imaging consisted of a dynamic phase of 2.5 h (first 10 min of 1 s frame^-1^, then 5 s frame^-1^ until 30 mins, then 10 s frame^-1^for the last 2 h) followed by a 10 min static scan. All images were processed and analysed using VivoQuant™ software (InVicro). A region of interest (ROI) was manually drawn around the heart and this area was applied to all images and timeframes generated in the scanning protocol.

### Least squares fitting

The time activity curves calculated at times t_1_, t_2_, …, t_N_ were normalised to the peak activity and fit simultaneously to the differential equations Eqs. (2-6), using lsqcurvefit least squares curve fitting and the ODE solver ode23s in MATLAB (MathWorks^**®**^). All 12 variables *k*_*f*_, *k*_1_, *E*_*m*_, *k*_2_, ΔΨ_*m*_, *t*_0_, *α, β, A*_0_, *V*_*p*_, *V*_*t*_ and *B* were allowed to vary freely within given boundary conditions. The values of *E*_*m*_ and ΔΨ_*m*_ were defined as positive voltages in the fitting routine so that all variables were positive. Errors in the least squares fitting were assessed by calculating the coefficient of variance, the covariance, correlation matrix and Jacobian returned from the lsqcurvefit function. Least squares fitting was used to derive initial parameter estimates for the MCMC procedure.

### Markov Chain Monte Carlo (MCMC) parameter estimation

The MCMC procedure used a Metropolis–Hastings random walk. All experimental data were analysed with 10,000 iterations. At each iteration of the procedure a new parameter estimate was calculated by adding a normally distributed random number multiplied by a scaling factor that defines the sensitivity of the random walk. The new parameters were accepted or rejected based on a Log likelihood function assuming fixed noise variance. An acceptance rate was calculated and the scaling factor of the proposed parameter estimate was adaptively adjusted automatically to achieve a target acceptance rate of 0.25. The initial burn-in period was 1000 iterations which were discarded, and the remaining parameter estimates were plotted as a histogram and mean, and standard deviation were calculated to estimate the best fit parameters and errors associated.

## Abbreviations

KHB: Krebs Henseleit buffer
MCMC: Markov Chain Monte Carlo
PET: Positron Emission Tomography
SPECT: Single Photon Emission Computed Tomography.

## Data, Materials, and Software Availability

All study data are included in the article. MATLAB code for the fitting procedure is given in the SI Appendix.

## Acknowledgements

We thank Prof Philip W. Kuchel for critically reading the manuscript and for advising on the use of Markov Chain Monte Carlo methods for parameter estimation by systematic random sampling of high-dimensional probability distributions.

## Author contributions

E.C.T.W, M.J.S., R.S. and T.R.E designed research; E.C.T.W. and F.B performed experiments; M.R.O. and T.R.E developed theory and wrote the MATLAB code; T.R.E analyzed data; T.R.E wrote the paper. All authors gave comments and approved the final manuscript.

## References

1 Hodgkin AL, Huxley AF & Katz B. Measurement of current-voltage relations in the membrane of the giant axon of Loligo. J Physiol. 1952;116(4):424–448.

2 Hodgkin AL & Katz B. The effect of sodium ions on the electrical activity of giant axon of the squid. J Physiol. 1949;108(1):37–77.

3 Mitchell P. Coupling of phosphorylation to electron and hydrogen transfer by a chemi-osmotic type of mechanism. Nature. 1961;191:144–148.

4 Pieczenik SR & Neustadt J. Mitochondrial dysfunction and molecular pathways of disease. Exp Mol Pathol. 2007;83(1):84–92.

5 Snabaitis AK, Shattock MJ & Chambers DJ. Comparison of polarized and depolarized arrest in the isolated rat heart for long-term preservation. Circulation. 1997;96(9):3148–3156.

6 Yang M & Brackenbury WJ. Membrane potential and cancer progression. Front Physiol. 2013;4.

7 Kowaltowski AJ & Abdulkader F. How and when to measure mitochondrial inner membrane potentials. Biophys J. 2024.

8 Rottenberg H. Membrane potential and surface potential in mitochondria: uptake and binding of lipophilic cations. J Membr Biol. 1984;81(2):127–138.

9 Kamo N, Muratsugu M, Hongoh R & Kobatake Y. Membrane potential of mitochondria measured with an electrode sensitive to tetraphenyl phosphonium and relationship between proton electrochemical potential and phosphorylation potential in steady state. J Membr Biol. 1979;49(2):105–121.

10 Kauppinen R. Proton Electrochemical Potential of the Inner Mitochondrial-Membrane in Isolated Perfused Rat Hearts, as Measured by Exogenous Probes. Biochim Biophys Acta. 1983;725(1):131–137.

11 Wan B, Doumen C, Duszynski J, Salama G, Vary TC & LaNoue KF. Effects of cardiac work on electrical potential gradient across mitochondrial membrane in perfused rat hearts. Am J Physiol. 1993;265(2 Pt 2):H453–460.

12 Fukuda H, Syrota A, Charbonneau P, Vallois J, Crouzel M, Prenant C et al. Use of 11C-triphenylmethylphosphonium for the evaluation of membrane potential in the heart by positron-emission tomography. Eur J Nucl Med. 1986;11(12):478–483.

13 Gurm GS, Danik SB, Shoup TM, Weise S, Takahashi K, Laferrier S et al. 4-[18F]-tetraphenylphosphonium as a PET tracer for myocardial mitochondrial membrane potential. JACC Cardiovasc Imaging. 2012;5(3):285–292.

14 Momcilovic M, Jones A, Bailey ST, Waldmann CM, Li R, Lee JT et al. In vivo imaging of mitochondrial membrane potential in non-small-cell lung cancer. Nature. 2019;575(7782):380–384.

15 Alpert NM, Guehl N, Ptaszek L, Pelletier-Galarneau M, Ruskin J, Mansour MC et al. Quantitative in vivo mapping of myocardial mitochondrial membrane potential. PLoS One. 2018;13(1):e0190968.

16 Pelletier-Galarneau M, Petibon Y, Ma C, Han P, Kim SJW, Detmer FJ et al. In vivo quantitative mapping of human mitochondrial cardiac membrane potential: a feasibility study. Eur J Nucl Med Mol Imaging. 2021;48(2):414–420.

17 Kawamoto A, Kato T, Shioi T, Okuda J, Kawashima T, Tamaki Y et al. Measurement of technetium-99m sestamibi signals in rats administered a mitochondrial uncoupler and in a rat model of heart failure. PLoS One. 2015;10(1):e0117091.

18 Safee ZM, Baark F, Waters ECT, Veronese M, Pell VR, Clark JE et al. Detection of anthracycline-induced cardiotoxicity using perfusion-corrected Tc sestamibi SPECT. Sci Rep. 2019;9.

19 Kuchel PW, Naumann C, Puckeridge M, Chapman BE & Szekely D. Relaxation times of spin states of all ranks and orders of quadrupolar nuclei estimated from NMR z-spectra: Markov chain Monte Carlo analysis applied to Li-7(+) and Na-23(+) in stretched hydrogels. J Magn Reson. 2011;212(1):40–46.

20 Veech RL, Kashiwaya Y, Gates DN, King MT & Clarke K. The energetics of ion distribution: the origin of the resting electric potential of cells. IUBMB Life. 2002;54(5):241–252.

21 Renkin EM. Transport of potassium-42 from blood to tissue in isolated mammalian skeletal muscles. Am J Physiol. 1959;197:1205–1210.

22 Crone C. The Permeability of Capillaries in Various Organs as Determined by Use of the ‘Indicator Diffusion’ Method. Acta Physiol Scand. 1963;58:292–305.

23 Kontos MC, Jesse RL, Schmidt KL, Ornato JP & Tatum JL. Value of acute rest sestamibi perfusion imaging for evaluation of patients admitted to the emergency department with chest pain. J Am Coll Cardiol. 1997;30(4):976–982.

24 Maddahi J & Packard RR. Cardiac PET perfusion tracers: current status and future directions. Semin Nucl Med. 2014;44(5):333–343.

25 Hershgold EJ, Steiner SH & Sapirstein LA. Distribution of myocardial blood flow in the rat. Circ Res. 1959;7(4):551–554.

26 Michaels AM, Zoccarato A, Hoare Z, Firth G, Chung YJ, Kuchel PW et al. Disrupting Na(+) ion homeostasis and Na(+)/K(+) ATPase activity in breast cancer cells directly modulates glycolysis in vitro and in vivo. Cancer Metab. 2024;12(1):15.

27 Lynch RM & Balaban RS. Coupling of aerobic glycolysis and Na+-K+-ATPase in renal cell line MDCK. Am J Physiol. 1987;253(2 Pt 1):C269–276.

28 Okamoto K, Wang W, Rounds J, Chambers EA & Jacobs DO. ATP from glycolysis is required for normal sodium homeostasis in resting fast-twitch rodent skeletal muscle. Am J Physiol Endocrinol Metab. 2001;281(3):E479–E488.

29 Cherry SR, Badawi RD, Karp JS, Moses WW, Price P & Jones T. Total-body imaging: Transforming the role of positron emission tomography. Sci Transl Med. 2017;9(381).

30 Cherry SR, Jones T, Karp JS, Qi J, Moses WW & Badawi RD. Total-Body PET: Maximizing Sensitivity to Create New Opportunities for Clinical Research and Patient Care. J Nucl Med. 2018;59(1):3–12.

31 Handley MG, Medina RA, Mariotti E, Kenny GD, Shaw KP, Yan R et al. Cardiac Hypoxia Imaging: Second-Generation Analogues of Cu-ATSM. J Nucl Med. 2014;55(3):488–494.

32 Medina RA, Mariotti E, Pavlovic D, Shaw KP, Eykyn TR, Blower PJ et al. Cu-CTS: A Promising Radiopharmaceutical for the Identification of Low-Grade Cardiac Hypoxia by PET. J Nucl Med. 2015;56(6):921–926.

