## Supplementary Information for "Sarcolemmal and mitochondrial membrane potentials measured *ex vivo* and *in vivo* in the heart by pharmacokinetic modelling of [^99m^Tc]sestamibi"

### Matlab code

```
% A MATLAB script to perform a Markov Chain Monte Carlo (MCMC) fit to time
% activity curve of [99mTc]sestamibi in vivo following a bolus injection
%
% The program performs an initial least squares fitting to optimize the
% input vector of parameters which are then used for teh MCMC estimation
%
% Authors: T.R. Eykyn, M.R. Orton
%

function [final_params] = Tc_fitting_TE_monteMCMCinvivo(fname, num_samples, burn_in)

    % Load and normalise data
    [t,dataH] = textread(fname,'%f %f');
    data = dataH / max(dataH);

    % Shift time scale so bolus arrival time is between 0s and 5s
    t = t - 57;
    data(t < 0) = [];
    t(t < 0) = [];

    % Initialise parameter vector and bounds
    %xIn =  [ kf   k1    Es     k2    Em    t0   a   b  A0  V1   V2   B    ]
    xIn  =  [ 6.2  0.2   0.065  0.03  0.15  2.6  14  5  1   0.4  0.3  0.01 ];
    r = 1 + 0.01 * randn(1, 12);
    xIn  = xIn .* r;
    lb = xIn/10;
    ub = 10*xIn;

    % Set optimizer options and perform least squares fit to the function Predict_invivo
    options=optimset('MaxIter',1e4,'MaxFunEvals',1e4,'TolFun',1e-4);
    xOpt = lsqcurvefit(@Predict_invivo,xIn,t,data,lb,ub,options);

    % calculate residuals and jacobian of the best fit
    [p_est,resnorm,residual,~,~,~,J] = lsqcurvefit(@Predict_invivo,xOpt,t,data,[],[],optimset('Display','off','MaxFunEvals',1));

    % Calculation of the coefficient of variance for error estimation
    gamma = resnorm/(length(data) - length(p_est));
    pCov = gamma*inv(J'*J);
    pSD = sqrt(diag(pCov))';
    pCV = 100*pSD./abs(p_est);
    pCorr = abs(pCov./sqrt(diag(pCov)*diag(pCov)'));

    % Display the values of the unknown parameters resulting from the least
    % squares fitting
    fprintf('  kf     k1     Es     k2     Em     t0     a     b     A0    V1    V2     B  \n')
    fprintf(' %.3f  %.3f  %.3f  %.3f  %.3f  %.2f  %.2f  %.2f  %.2f  %.2f  %.2f  %.3f  \n',xOpt)
```

```
    % Plot the fit for the final set of parameters with residuals and correlation matrix
    [M,M1,M2,M3] = Predict_invivo(xOpt,t);
    figure; imagesc(pCorr);
    figure; subplot(1,2,1); semilogx(t,data,'k.',t,M(1:length(t)),'b',t,M1,'k',t,M2,'r',t,M3,'g','Linewidth',1);
    axis tight; axis square; xlim([1 max(t)]); ylim([0 1]);
    subplot(1,2,2); semilogx(t,residual(1:length(t)),'.k'); axis tight; axis square; xlim([1 max(t)]); ylim([0 1])

    % If least squares gives a good fit then continue to perform MCMC estimation
    prompt = "Do you want to run a Monte Carlo fitting y/n?";
    txt = input(prompt,"s");

    if ismember('y',txt)
        close all;
        xIn = xOpt;
        lb = xIn/5;
        ub = 5*xIn;
        [samples] = MCMC(t, data, xIn, lb, ub, num_samples, burn_in);

        % Discard burn-in period
        final_params.lsq = xOpt;
        final_params.all = samples(burn_in+1:end,:);
        final_params.mean = mean(samples(burn_in+1:end,:));
        final_params.std = std(samples(burn_in+1:end,:));

        % Plot the fit for the final set of parameters
        [M, M1, M2, M3] = Predict_invivo(final_params.mean, t);
        figure;
        semilogx(t, data, 'k.', t, M, 'b', t, M1, 'k', t, M2, 'r', t, M3, 'g', 'Linewidth', 1);
        axis tight; axis square; xlim([1 max(t)]);
        figure; plot(final_params.all(1:end,3)); hold on; plot(final_params.all(1:end,5)); axis tight;
        figure; histogram(final_params.all(:,3),20); hold on; histogram(final_params.all(:,5),20); xlim([0 0.2]);
    else
        final_params.lsq = xOpt;
        close all;
    end
end

function [samples] = MCMC(t, data, xIn, lb, ub, num_samples, burn_in);

    % Initialize MCMC parameters
    current_params = xIn;
    current_likelihood = likelihood(data, t, current_params);

    % Store samples and acceptance rate
    samples = zeros(num_samples, length(xIn));
    accepted = 0;
    proposal_scale = 1e-3; % Initial proposal scale, change this number to adjust sensitivity of MCMC

    % MCMC Sampling loop
```

```
    for i = 1:num_samples
        % Propose new parameters using a Gaussian distribution
        proposed_params = current_params + proposal_scale*randn(1, length(xIn));

        % Vectorized bounds check
        if all(proposed_params >= lb) && all(proposed_params <= ub)
            % Calculate likelihood of proposed parameters
            proposed_likelihood = likelihood(data, t, proposed_params);

            % Metropolis-Hastings acceptance criterion (log space to avoid numerical issues)
            log_acceptance_ratio = proposed_likelihood - current_likelihood;

            if log(rand()) < log_acceptance_ratio
                current_params = proposed_params;
                current_likelihood = proposed_likelihood;
                accepted = accepted + 1;
            end
        end

        % Store the current parameters as a sample
        samples(i, :) = current_params;

        % Display progress (optional)
        if mod(i, 200) == 0
            acceptance_rate = accepted / i;
            fprintf('Iteration %d/%d - Acceptance rate: %.2f - Proposal Scale: %.2e - Current likelihood: %.2e\n',...
                i, num_samples, acceptance_rate, proposal_scale, current_likelihood);
        end

        % Plot parameter evolution (optional)
        if mod(i, 200) == 0
            plot(samples(1:i,3)); hold on; plot(samples(1:i,5)); axis tight;
            drawnow;
        end

        % Adjust proposal scale adaptively every 100 iterations to achieve
        % a given target acceptance
        target_acceptance = 0.25;
        if mod(i, 100) == 0
            acceptance_rate = accepted / i;
            adjustment_factor = exp((acceptance_rate - target_acceptance) * 0.1);
            proposal_scale = proposal_scale * adjustment_factor;
        end
    end
end

function logL = likelihood(data, t, params)
    [M, ~, ~, ~] = Predict_invivo(params, t);
    residual = data - M;
```

```
    sigma2 = var(data);
    logL = -0.5 * sum((residual.^2) / sigma2 + log(2 * pi * sigma2));
end

function [M, M1, M2, M3] = Predict_invivo(xIn, tIn)
    [~, y] = ode23s(@rigid_invivo, tIn, [0,0,0], [], xIn);
    V1  = xIn(10);
    V2  = xIn(11);
    V3 = 1 - V1 - V2;
    B = xIn(12);
    M1 = V1 * y(:,1);
    M2 = V2 * y(:,2);
    M3 = V3 * y(:,3);
    M = M1 + M2 + M3 + B;
end

function dy = rigid_invivo(t, y, xIn)
    dy = zeros(3, 1);
    R = 8.314;
    T = 310;
    F = 96485;

    kf  = xIn(1);
    k1  = xIn(2);
    k_1 = xIn(2) * exp((-xIn(3) * F) / (R * T));
    k2  = xIn(4);
    k_2 = xIn(4) * exp((-xIn(5) * F) / (R * T));

    kloss = 3.2059e-05;
    U = ArterialInput(t - xIn(6), xIn(7), xIn(8), xIn(9));

    dy(1) =  kf * U - (kf - k1 + kloss) * y(1) + k_1 * y(2);
    dy(2) =  k1 * y(1) - (k_1 + k2 + kloss) * y(2) + k_2 * y(3);
    dy(3) =  k2 * y(2) - (k_2 + kloss) * y(3);
end

function [c] = ArterialInput(tH, a, b, A0)
    c = A0 * (tH.^a) .* exp(-tH * b);
    c(tH < 0) = 0;
end
```
